# The ferroptosis inhibitor NUPR1 coordinates the mitochondrial response to oxidative stress and cell metabolism during COPD pathogenesis in the lung

**DOI:** 10.1101/2025.04.27.650854

**Authors:** Saul S. Siller, Tao Yang, Jessica Nouws, So-Jin Kim, Sang-Hun Kim, Patricia Santofimia-Castaño, Juan Iovanna, John E. McDonough, Maor Sauler

## Abstract

Chronic obstructive pulmonary disease (COPD) is characterized by chronic injury and oxidative stress leading to progressive lung tissue destruction. Emerging evidence suggests that regulated cell death pathways, particularly ferroptosis, contribute to COPD pathology. We previously identified the stress response protein and known ferroptosis inhibitor nuclear protein 1 (NUPR1) as markedly downregulated in lung tissue from COPD patients. Here, we demonstrate that NUPR1 inhibition exacerbates iron accumulation, enhances lipid peroxidation, impairs mitochondrial function, disrupts cellular metabolism, and increases oxidative stress in lung epithelial cells. Furthermore, *Nupr1* knockout mice exhibit mitochondrial abnormalities, increased oxidative damage, and lung tissue changes consistent with COPD pathology. Collectively, our findings establish NUPR1 as a critical regulator of ferroptosis, stress responses, mitochondrial integrity, and metabolic balance in the lung, highlighting its potential contribution to COPD pathogenesis.

## Introduction

Chronic obstructive pulmonary disease (COPD) is a leading cause of morbidity and mortality in the United States (1). It is clinically defined by persistent airflow obstruction and is often accompanied by emphysema, a pathological manifestation characterized by chronic inflammation and alveolar destruction (2, 3). Recent findings suggest that multiple regulated cell death pathways, including ferroptosis, contribute to the development of COPD and emphysema (4–8). Indeed, cigarette smoke induces epithelial ferroptosis in the lung while inhibition of ferroptosis reduces emphysematous pathology in mouse models (9).

Ferroptosis is an iron-dependent regulated cell death pathway resulting from redox imbalance and characterized by excessive lipid peroxidation (10, 11). As a consequence of oxidative stress, excess ferrous iron (Fe^2+^) causes damage by converting hydrogen peroxide into highly reactive hydroxyl radicals, which promotes oxidization of membrane lipids (12). This lipid peroxidation undermines ion transport and causes cellular swelling that ultimately leads to rupture of the plasma membrane and cell death (13). Cells mitigate ferroptosis primarily by controlling iron homeostasis, maintaining adequate antioxidant defenses, and limiting excessive lipid peroxidation (14). One critical mechanism is the production of glutathione (GSH) from glutamate and cysteine, which requires cystine import via the cystine/glutamate antiporter system X_C_^-^ and subsequent conversion to cysteine (15–18). Glutathione peroxidase 4 (GPX4) then uses GSH to detoxify lipid hydroperoxides, preventing the propagation of oxidative damage to membranes (19). Consequently, the availability of cysteine and glutamate is critical for controlling oxidative stress. Additional protective pathways include iron chelation or sequestration to limit redox-active iron as well as other antioxidant systems, including the FSP1-CoQ_10_ axis and NRF2 signaling, which both support the cellular redox balance and prevent buildup of harmful lipid peroxides (10, 16). Finally, TP53, in coordination with distinct members of the ALOX family, can both promote and inhibit ferroptosis in a cell- and tissue-specific manner (20). Given these multiple layers of regulation and the importance of ferroptosis in COPD pathogenesis, it is crucial to identify the factors that affect an individual’s susceptibility to ferroptotic processes in the lung.

Recent studies have implicated nuclear protein 1 (NUPR1) in the regulation of ferroptosis (21). NUPR1 is an intrinsically disordered protein that can shuttle to the nucleus and function as a transcription co-regulator by interacting with other DNA-binding proteins and regulatory complexes (22, 23). Numerous reports suggest that NUPR1 influences mitochondrial biology, iron metabolism, and ferroptosis (24–27); however, its role in the lung remains largely unexplored. Previously, we found that alveolar type 2 (AT2) cells from subjects with emphysema exhibit marked downregulation of NUPR1 and that NUPR1 inhibition increases susceptibility to iron-dependent cell death following cigarette smoke extract (CSE) exposure (28). In the present work, we demonstrate that NUPR1 inhibition drives iron accumulation, increases lipid peroxidation, disrupts metabolism, impairs mitochondrial integrity and function, and elevates reactive oxygen species (ROS) generation. Collectively, these findings establish NUPR1 as a critical regulator of ferroptosis, mitochondrial homeostasis, and cellular metabolism in the lung.

## Materials and Methods

### Cell culture

A549 cells were cultured in DMEM supplemented with 10% FBS (R&D systems, S11150H) in 5% CO2 and at 37°C. Human bronchial epithelial cells (HBECs) were cultured in Pneumacult-Ex Medium (Stemcell Technologies, 5008) in 5% CO2 and at 37°C. For CSE experiments, 20 mL of cell culture media was bubbled, under negative pressure, with mainstream smoke from two 3RF4 research cigarette (University of Kentucky, Lexington, Kentucky, USA). The CSE was filtered via 0.22 μM filter (MilliporeSigma) to obtain a filtrate that was considered as 100% CSE. The CSE was then aliquoted and stored at −80°C.

### RNAi transfection

Cells were treated with RNAiMAX according to manufacturer’s protocol. Targets were purchased from Horizon. OnTARGETplus non-targeting pool (D-0001810-10-01) was used as a negative control (control), and OnTARGETplus Human NUPR1 (26471)- smartpool (L-012819-00-0005) was used for NUPR1 knockdown (KD). For all functional assays, 5×10^5^ A549 cells were collected after two 72 hour cycles of RNAi transfection unless otherwise noted.

### Cell ROX assay

Transfected A549 cells were treated with 5% CSE overnight, then harvested, washed in PBS, and incubated in Mitotracker red CMXRos (M7512) and Mitotracker green FM (M7514) for 45 minutes at 37°C in the dark. After this incubation, cells were washed 1X with PBS and analyzed by flow cytometry (CytoFLEX flow cytometer). Cells positive for red and green fluorescence were counted and divided. Control was normalized to 1.

### Quantification of mtDNA content

DNA was isolated with the Qiagen blood and tissue kit from harvested A549 cells. Subsequently, we performed a RT-qPCR using SYBR with primers specific for the mitochondrial tRNALeu^(UUR)^ gene and the nuclear β-2-microglobulin (β*2M*) as a reference gene (29). Primer sequences were as follows: mtDNA tRNALeu(UUR) tRNA F, CACCCAAGAACAGGGTTTGT; mtDNA tRNALeu(UUR) tRNA R, TGGCCATGGGTATGTTGTTA; nDNA β2-microglobulin ß2M F, TGCTGTCTCCATGTTTGATGTATCT; nDNA β2-microglobulin ß2M R, TCTCTGCTCCCCACCTCTAAGT. Relative mtDNA expression was calculated with the 2^-ΔΔ*CT*^ method, with control cells relative tRNALeu^(UUR)^ expression level being normalized to 1.

### Mitochondrial depolarization assay

A549 cells were processed according to the manual provided by the MitoProb JC-1 Assay Kit for Flow Cytometry (M34152). For positive control of mitochondrial depolarization, we treated cells with 10 mM carbonyl cyanide m-chlorophenyl hydrazone (CCCP) (ab141229) for 4 hrs prior to harvesting.

### Lipid peroxidation assays

Transfected cells were incubated with 10 μM erastin and 1 μM ferrostatin overnight with Bodipy dye (D3861) for 30 minutes at 37°C in a 12 well plate. Cells were harvested and washed 2x in PBS, and green fluorescence was detected by flow cytometry. Control was normalized to 1.

### 8-OHdG

Cultured A549 cells were washed in PBS and stained with 8-OHdG-AF647 (Santa Cruz, sc393871), for 1 hour at 4°C, and subsequently washed 2x in PBS before analyzing by flow cytometry. Cells positive for 8-OHdG were counted, and control was normalized to 1.

### Cell death assay

5×10^5^ HBECs were harvested after two 72 hour cycles of RNAi. Cells were washed 2X in PBS and incubated with Annexin-V-FITC and PI for 30 minutes at 37°C in annexin binding buffer. Cells were immediately analyzed by flow cytometry. Cell death is expressed as percentage of cells.

### Iron studies

Transfected A549 cells were incubated without or with 10 μM erastin overnight. Fe^2+^ levels were then assessed by the FerroOrange Intracellular Iron Measurement kit (Dojindo F374) as per the manufacturer’s protocol. Cellular fluorescence was then measured by flow cytometry, and control was normalized to 1.

### Cell viability/MTT assays

Cell viability was assayed using MTT (3-(4,5-Dimethylthiazol-2-yl)-2,5-Diphenyltetrazolium Bromide) (Roche #1465007) as previously described (28). In brief, A549 cells were incubated with erastin at noted concentrations and 1 µM ferrostatin overnight. Subsequently, cells were incubated overnight with MTT reagent in growth medium as per manufacturer’s protocol. Absorbance was then measured the following day at 550 nm, and control was normalized to 1.

### Transmission electron microscopy

Transmission electron microscopy (TEM) was performed as previously described (30). In brief, mouse lung tissue was cut in 2 mm^2^ blocks while A549 cells were grown and transfected in a 10 cm dish. The tissue and cells were fixed in 2.5% glutaraldehyde in 0.1 M sodium cacodylate buffer, pH 7.4, for 1 hr and then rinsed with sodium cacodylate buffer, scraped, and pelleted in 2% agar. Samples were trimmed and post-fixed with 1% osmium tetroxide for 1 hr. After being stained in 2% uranyl acetate in maleate buffer, pH 5.2, for 1 hr, samples were dehydrated in an ethanol series, infiltrated with resin (Embed812, Electron Microscopy Science), and allowed to cure overnight at 60°C. Hardened blocks were then sliced with a Leica UC7 Ultramicrotome, and 60 nm sections were collected onto formvar/carbon coated nickel grids. These sections were subsequently stained with 2% uranyl acetate and lead citrate and viewed with a FEI Tecnai Biotwin TEM at 80 kV. Images were taken using Morada CCD and iTEM (Olympus) software typically at 26,000x magnification.

### Mouse models and husbandry

*Nupr1*^-/-^ mice were obtained from Juan Iovanna (CRCM, France) (31). Mice were bred and maintained according to protocols approved by the Institutional Animal Care and Use Committee at Yale University (IACUC protocol 07867). Primer sequences for genotyping were as follows: NUPR1^+/+^ F, CAGCCCATCTGCTTCTCACT; NUPR1^+/+^ R, GGGTTACTTGGGAGTTGGGAATA; NUPR1^-/-^ F, GTCAGCCCATCTGCTTCTCA; NUPR1^-/-^ R, TGTTTTGCCAAGTTCTAATTCCATCAGA.

### Immunostaining and imaging

Mouse lungs were fixed in formalin for 24 hours, dehydrated in 70% ethanol, and then embedded in paraffin. The paraffin embedded blocks of mouse lung tissue samples were deparaffinized in xylene and decreasing concentrations of ethanol in distilled water. They were then placed in citrate, pH 6, epitope retrieval buffer (Biolegend, 420901) at 95°C for 20 min, cooled, and permeabilized in PBS with 0.1% Tween (PBSTw) and 0.2% Triton-X (PBST) for 10 minutes. The samples were subsequently rinsed and washed in PBSTw. Slides were incubated in CAS-Block (Life Technologies, 008120) for 10 minutes. Primary antibodies were applied overnight at 4°C and included the following: rabbit anti-SPC (Millipore#AB3786, 1:200), mouse anti-SOD2 (Santa Cruz sc#137254, 1:50), and mouse 8-OHdG (Santa Cruz #sc-66036, 1:50). Slides were washed with PBSTw and then incubated for 1 hour at room temperature with the following secondary antibodies: donkey anti-mouse Alexa-488 (Invitrogen) and donkey anti-rabbit Alexa-568 (Invitrogen). Slides were then again washed with PBSTw. Coverslips were mounted using anti-fade mounting media with DAPI (Vectashield). Images were acquired with Nikon Eclipse Ti.

### Histology and cord length measurement

Hematoxylin and eosin (H&E) staining was performed on paraffin-embedded lung slices using standard protocols. Cord length measurements were conducted with ImageJ as previously described (32).

### Transcriptomic sample preparation, sequencing, and data analysis

5×10^5^ A549 cells were harvested after two rounds of RNAi. RNA was isolated following the manufacturer’s protocol of the RNeasy Mini Kit (Qiagen #74104) and then subjected to DNase treatment. Samples were then submitted to the Yale Center for Genome Analysis (YCGA) for poly(A)-selected library creation utilizing the Kapa Stranded RNA-Seq kit (Roche 07962193001). Sequencies was then performed by the YCGA on a NovaSeq6000 platform. RNA-sequencing data was then processed using standard methods with a log_2_(fold change) cut-off of ≤-0.4 or ≥0.4. For Gene Set Enrichment Analysis (GSEA), publicly available software (https://www.gsea-msigdb.org/gsea/index.jsp) was employed as described previously (33, 34).

### METAFlux data analysis

METAFlux software was employed as previously described (35), utilizing standard features and parameters with bulk RNA sequencing data as input.

### Metabolomic sample preparation, mass spectrometry, and data analysis

For sample preparation, cells were cultured in a 6-well plate and quickly washed 1X with ice cold 5 mM HEPES (2 mL/well). They were then harvested on ice with ice cold quench buffer (150 µL/well, composed of 20% methanol, 3 mM sodium fluoride, 32 µM D_8_-phenylalanine as internal standard and 0.1% formic acid). Cell lysates lyophilized in a 96-well plate. Samples were prepared by resuspending the cell pellet in 50 µL 10% acetonitrile solution with D_4_-taurine (25 µM) as a second internal standard. 5 µL of the supernatant was injected for each analysis mode on the mass spectrometer.

For chromatography, two columns were used separately, a Thermo Scientific Hypercarb Porous Graphitic Carbon HPLC Column (100 x 4.6 mm, 3 µm) and a Phenomenex Kinetex F5 Core-shell HPLC column (100 x 2.1 mm, 2.6 µm). Separation by the Hypercarb column was performed with 1 mL/min linear gradients as indicated (Supplementary Table 1). Column temperature was maintained at 50°C with the auto-sampler at 5°C. Separation by the Kinetex F5 column was performed with 0.3 mL/min linear gradients as outlined (Supplementary Table 2). Column temperature was maintained at 30°C with the auto-sampler again at 5°C.

For the mass spectrometry, data were analyzed using the Sciex TripleTOF 6600 collected using an information-dependent analysis (IDA) workflow, consisting of a TOF MS scan (200 msec) and a high-resolution IDA experiment (70 msec each) monitoring 10 candidate ions per cycle. Former target ions were excluded after 2 occurrences for 5 sec and dynamic background subtraction was employed. The mass range for both TOF MS and IDA MS/MS scans was 60-1000 with the RP column and 70-1000 with the Hypercarb column.

The ion source conditions were as follows: ion spray voltage of 5000 V for the positive mode and −4500 V for the negative mode, ion source gas 1 (GS1) of 50, ion source gas 2 (GS2) of 50, curtain gas (CUR) of 30, and temperature of 400°C with the F5 column while 500°C with the Hypercarb column. Compound dependent parameters for all four modes were as follows: declustering potential (DP) of 35, collision energy (CE) of 30, and collision energy spread (CES) of 20.

For the data processing, El-MAVEN software (Elucidata.io) was used for peak picking and curation from house built targeted and untargeted libraries. Targeted libraries used the commercial standard kit of ∼600 metabolites (IROA Technologies) in all 4 modes of analyses (2 LC systems, positive and negative ion acquisition modes), yielding reference data (molecular ions and retention times) with wide coverage of the endogenous metabolome. Untargeted libraries contain ∼2,700 metabolites drawn KEGG database, and the top 5 candidates with the highest intensity throughout a sample run for each metabolite were automatically curated.

Data were normalized by the sum of all targeted metabolites within each sample in each mode and then log transformed. Sample quality was assessed using two internal standards, D_8_-phenylalanine and D_4_-taurine, to identify any outliers to be excluded from the downstream analysis. Next, the normalized data from all four modes were merged. If a metabolite appeared in more than one mode, the one with the best intensity was chosen.

PCA, PLS-DA and heat map on the merged targeted data were performed using MetaboAnalyst 5.0 (https://www.metaboanalyst.ca/). Differential expression analysis was performed using the metabolomics application in Polly (polly.elucidata.io). Normal p values and log2FC data was uploaded in Shiny GATOM (https://artyomovlab.wustl.edu/shiny/gatom/) to generate an optimized metabolic network displaying ∼60 of the most altered and closely connected metabolites. The up- and down-regulated metabolites in the NUPR1 KD group were then used separately for pathway enrichment analysis in MetaboAnalyst 5.0 (https://www.metaboanalyst.ca/).

### Statistics

Statistical methods, replicates, and p-values relevant to each figure are outlined in the figure legend. All statistical analysis was performed using GraphPad Prism 10 (GraphPad Prism Software). All error bars represent data±SEM.

## Results

### NUPR1 inhibits ferroptosis in the AT2 cells of the lung

We previously showed that NUPR1 depletion (via small interfering RNA [siRNA]) increases susceptibility to iron-dependent cell death after CSE exposure in the immortalized AT2 A549 cell line (28). To confirm that loss of NUPR1 specifically promotes ferroptosis, we treated A549 cells transfected with either a siRNA pool that were non-targeting or targeted to NUPR1 (control or NUPR1 KD, respectively) with erastin, a ferroptosis inducer (Fig. 1A) (15, 36). We found that cell viability NUPR1 KD was reduced to 54% compared to controls (p=0.0324 for control with erastin 10 μM vs NUPR1 KD with erastin 10 μM). Of import, with increasing doses of erastin, NUPR1 KD cell viability was diminished in a dose-dependent manner while erastin did not affect cell viability in control cells. Ferrostatin-1, a ferroptosis and lipid peroxidation inhibitor (36), rescued cell viability in NUPR1 KD cells. Consistent with a role in regulating ferroptosis and its upstream processes, we also observed increased erastin-induced lipid peroxidation (Fig. 1B, p<0.0001) and Fe2+ accumulation (Fig. 1C, p=0.0001) in NUPR1 KD cells. Next, we assessed whether NUPR1 depletion promotes ferroptosis in primary HBECs (Fig. 1D). CSE significantly increased cell death in NUPR1 KD HBECs (50.6% vs. 7.9%, p<0.0001), with partial rescue by the iron chelator deferoxamine (38.9%). Notably, deferoxamine did not affect CSE-induced cell death in controls (27.4 vs. 26.3% in CSE vs. CSE with deferoxamine controls). Collectively, these findings demonstrate a critical role for NUPR1 in regulating susceptibility to ferroptosis in respiratory epithelial cells.

**Figure 1.**
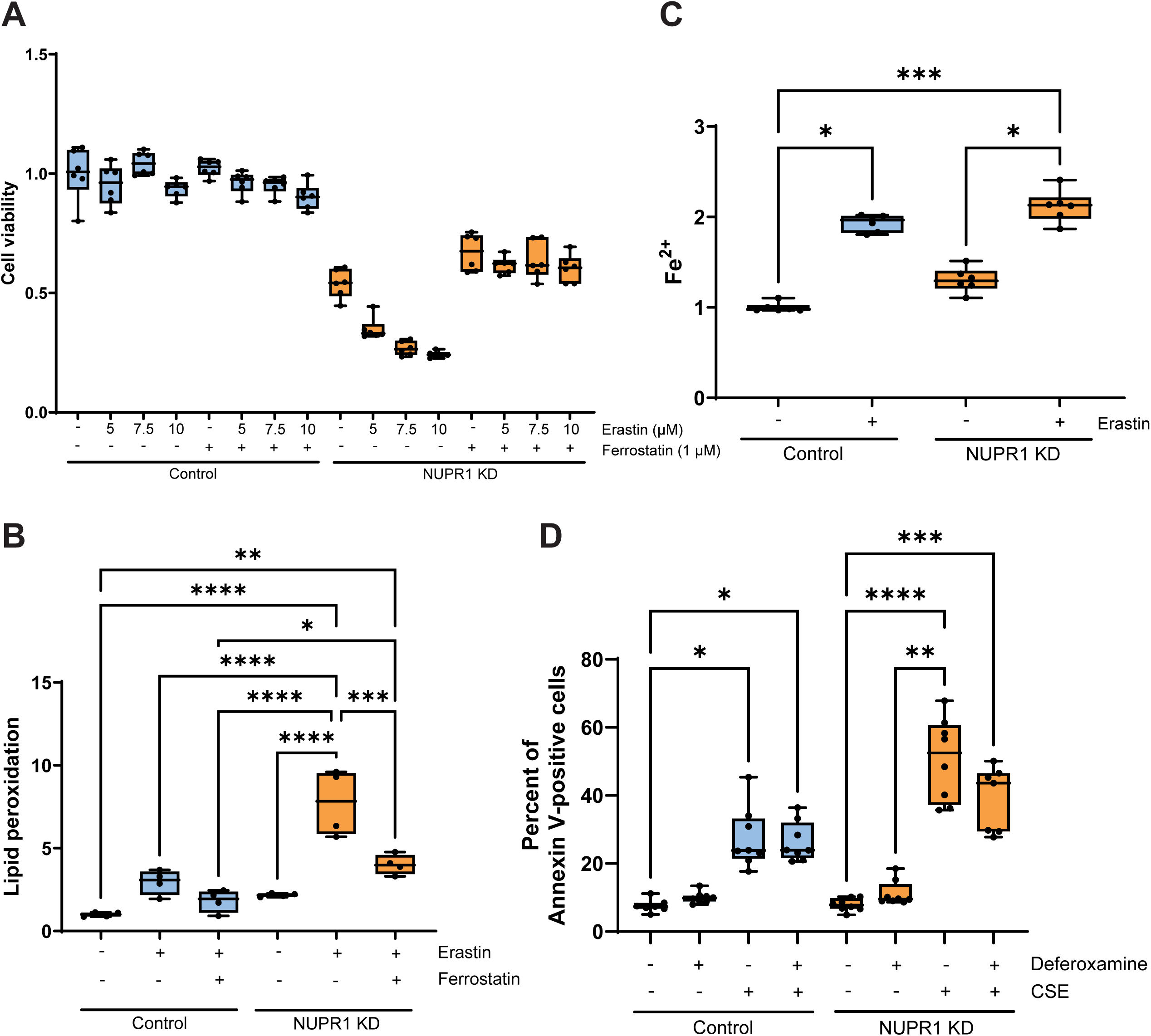
NUPR1 knockdown results in increased susceptibility to ferroptosis. (A) Cell viability assays are depicted with and without NUPR1 after treatment with vehicle, erastin, or ferrostatin. (B) Quantification of lipid peroxidation is shown for control and NUPR1 KD without or with erastin and/or ferrostatin. (C) Changes in Fe^2+^ concentration are observed for control and NUPR1 KD without or with erastin. (D) Annexin-V positivity was assessed for HBECs with or without NUPR1. CSE and/or 100 μM deferoxamine was present as indicated. Doses for pharmacologic treatment are indicated on all plots. Each point represents a replicate on graphs. One-way ANOVA tests were performed for all data with subsequent Kruskal-Wallis multiple comparison tests shown on graphs. *, p<0.05; **, p<0.01; ***, p<0.001; ***, p<0.0001.

### NUPR1 modulates the oxidative stress response and mitochondrial function in alveolar epithelial cells

The redox state of the cell influences the susceptibility to lipid peroxidation and ferroptosis (10). Indeed, we observe increased cellular redox after NUPR1 loss that is comparable to CSE of control and NUPR1 KD A549 cells (Fig. 2A, p<0.0001) (Figs. 2A). We also identified increased oxidative DNA damage in NUPR1 KD A549 cells as assessed by 8-hydroxy-2-deoxyguanosine (8-OHdG) (Fig. 2B, p=0.0392).

**Figure 2.**
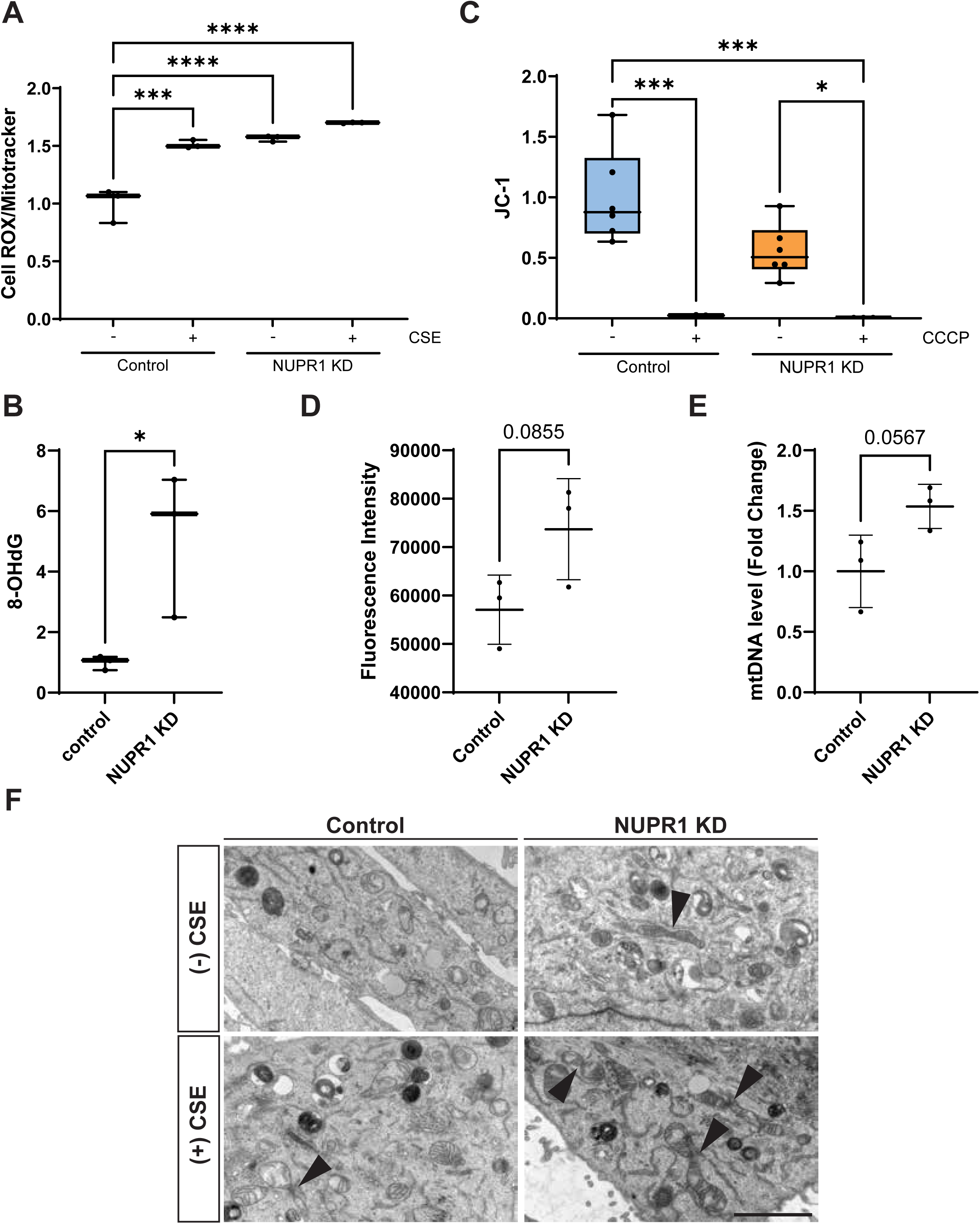
NUPR1 influences cellular redox, oxidative stress, and mitochondrial function. Graphs demonstrate levels of cellular redox normalized based on mitochondrial content without or with CSE (A), 8-OHdG (B), and mitochondrial membrane potential without or with CCCP (C) in control or NUPR1 KD A549 cells. Additional plots show quantification of mitochondrial content based on Mitotracker fluorescence intensity as assessed by flow cytometry (D) or mitochondrial DNA based on qPCR (E) in control and NUPR1 KD A549 cells is shown. Each point represents a replicate on graphs. One-way ANOVA tests were performed for redox and mitochondrial membrane potential data with subsequent Kruskal-Wallis multiple comparison tests shown on graphs. A two-tailed T-test was performed for the 8-OHdG and mitochondrial content data. *, p<0.05; ***, p<0.001; ***, p<0.0001. (F) TEM images of A549 cells with or without NUPR1 and/or CSE are depicted as indicated. Mitochondrial fission/fusion events are noted with arrowheads and observed with increasing frequency in NUPR1 KD cells and upon exposure to CSE. Scale bar, 2 μm.

Increased cellular redox and oxidative damage often result from dysfunctional mitochondria (37). To determine the impact of NUPR1 KD on mitochondrial function, we examined mitochondrial membrane potential and identified a reduction in mitochondrial membrane potential in NUPR1 KD compared to control A549 cells (Fig. 2C, p=0.0518). Interestingly, this reduction in membrane potential was observed in spite of an increase in mitochondrial content as assessed by flow cytometry and the amount of mitochondrial DNA (Fig. 2D-E). Finally, we examined CSE and non-CSE exposed A549 control or NUPR1 KD cells by TEM (Fig. 2F). Specifically, we observed an increased presence of mitochondrial fission/fusion events in TEM images of CSE-exposed compared to non-CSE exposed cells (68% vs. 50%). These fission/fusion events had a further increased presence in CSE-exposed NUPR1 KD cells (75%). Taken together, these findings suggest a critical role for NUPR1 in the regulation of cellular redox and mitochondrial function.

### Transcriptomic analysis reveals important functions for NUPR1 in iron metabolism, ferroptosis, and mitochondrial biology

To further delineate the function of NUPR1 in the lung, we performed RNA-sequencing of A549 cells with or without NUPR1 (Fig. 3A). We first confirmed a significant reduction in NUPR1 expression upon siRNA KD in A549 cells. After setting a threshold of a log_2_(fold change) ≤-0.4 or ≥0.4, we identified increased expression of genes associated with iron metabolism and ferroptosis (*LCN2*, *STEAP3, GPX4*) while the cystine/glutamate antiporter *SLC7A11* had decreased expression. We also noted increases in expression of certain key regulators of mitochondrial maintenance and dynamics (*LONP1*, *PINK1*, *MFN2*, *DNM2*) with decreases in others (*TFAM*, *MFN1*, *OPA1*). Quite intriguingly, multiple genes encoding subunits of the electron transport chain were all upregulated (*NDUFAB1*, *NDUFA4*, *NDUFS8*, *UQCRQ*, *UQCRB*, *COX8A*, *COX6B1*, *COX5B*, *COX7C*, *COX6C*, *ATP5F1D*, *ATP5F1E*). We additionally examined gene expression by RNA-sequencing of A549 cells with or without NUPR1 exposed to CSE utilizing the same log_2_(fold change) cut-offs. Many of these observed changes in gene expression persisted upon CSE exposure, albeit to a lesser degree. In agreement with these findings, gene set enrichment analysis (GSEA) further revealed disruptions in metabolic and cell death pathways (Fig. 3B). Taken together, these results underscore the critical regulation of ferroptosis, mitochondrial integrity, and metabolic pathways by NUPR1 in lung epithelial cells.

**Figure 3.**
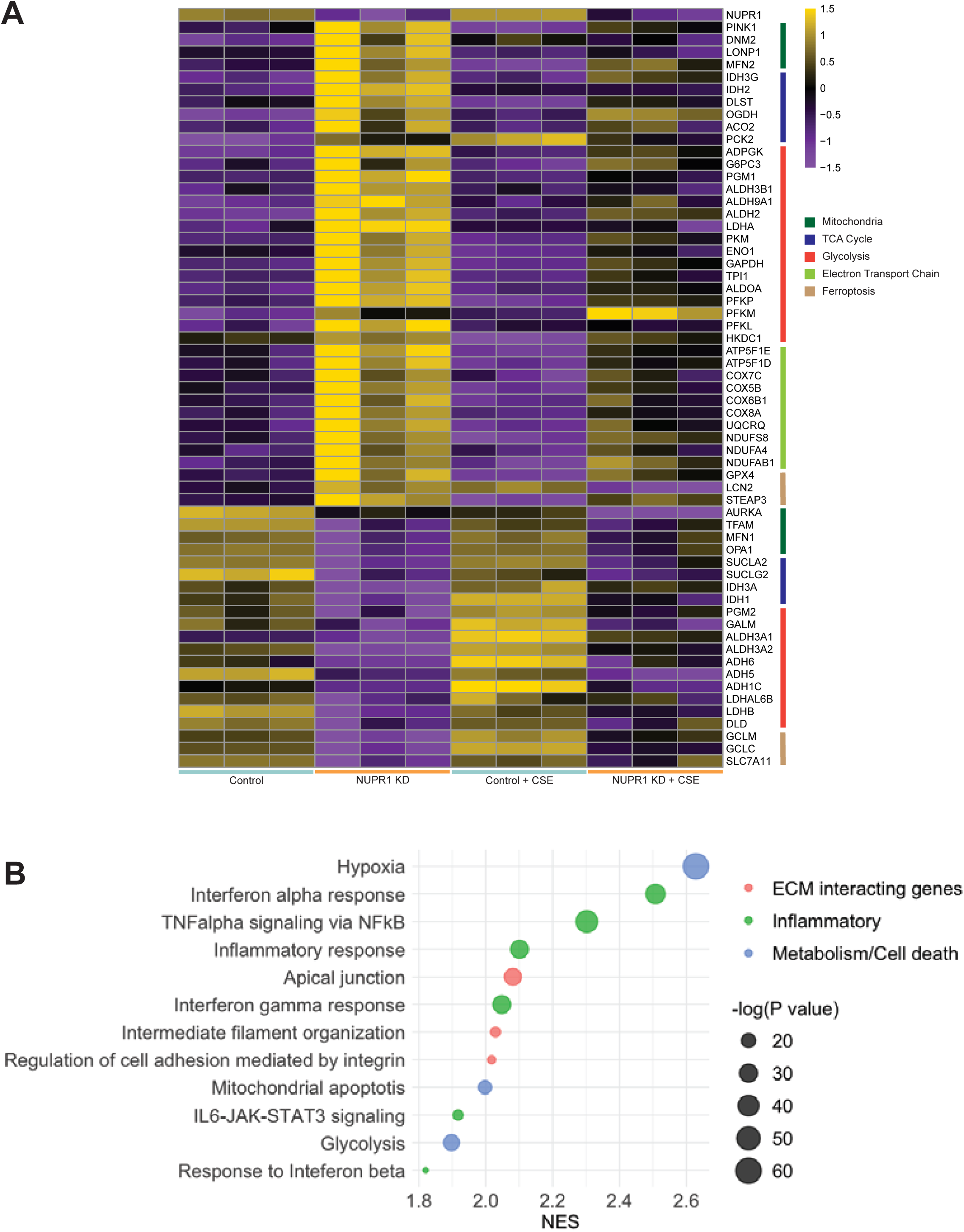
Transcriptomic analysis reveals significant expression changes in genes involved in iron metabolism, ferroptosis, mitochondrial function, and bioenergetics after NUPR1 loss. (A) Heatmap demonstrates up-regulation or down-regulation of expression for the selected genes. RNA-sequencing was performed in triplicate on A549 cells for the following conditions: control, NUPR1 KD, control + CSE, and NUPR1 KD + CSE as indicated. Expression thresholds of log_2_(fold change)≤-0.4 or ≥0.4 were utilized. All significant changes are shown with indicated color change based on scale representing log_2_(fold change). (B) Dot plot highlights multiple significantly enriched gene pathways based on GSEA using transcriptomic data as input. Size of dot corresponds to −log_10_(p value) as indicated while normalized enrichment score (NES) is shown on x-axis.

### NUPR1 substantially modulates cell metabolism in the lung

Our RNA-sequencing data revealed altered expression of mitochondrial and electron transport chain genes, and our GSEA highlighted disruptions in metabolic pathways. To further probe the function of NUPR1 in metabolism, we analyzed our RNA-sequencing results with METAFlux, which predicts the flux of metabolic reactions based on transcriptomic data and a defined metabolic milieu (35). METAFlux identified significant dysregulation of multiple pathways in A549 NUPR1 KD cells (Fig. 4A). Notably, alanine (Ala), aspartate (Asp), and glutamate (Glu) metabolism was significantly increased, congruent with an increased reliance on glutamate for system X_C_^-^ anti-transport of cystine and glutamate to fuel replenishment of GSH (13, 16–18, 38). Glycolysis/gluconeogenesis and oxidative phosphorylation also exhibited substantial changes in their fluxes, indicating that loss of NUPR1 significantly disrupts the metabolic landscape in A549 cells.

**Figure 4.**
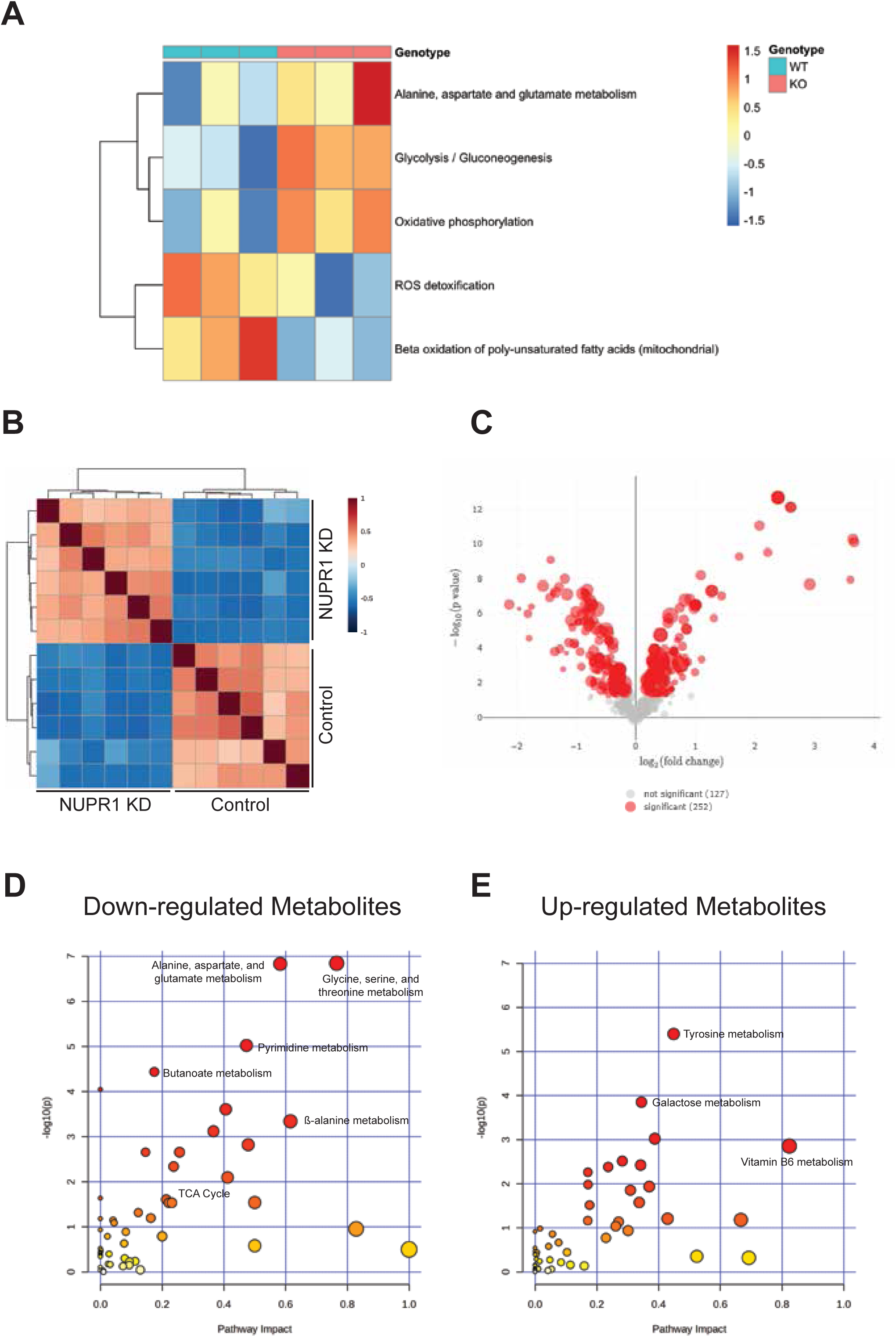
NUPR1 regulates multiple metabolic pathways related to antioxidant systems, glycolysis, the TCA cycle, and oxidative phosphorylation. (A) Heatmap displays altered flux in metabolic pathways for control or NUPR1 KD A549 cells. Changes in METAFlux scores utilizing A549 transcriptome data as input are represented by color changes on heatmaps. Only significant changes based on Wilcoxon rank sum analysis are shown. For metabolomic data, correlation analysis by sample (B) is shown. (C) Volcano plot demonstrates altered metabolites in NUPR1 KD versus control A549 cells based on log_2_(fold change) on the x-axis and on −log_10_(p value) on the y-axis. Plots showing results from pathway enrichment analysis for down-regulated (D) and up-regulated (E) metabolic pathways with the pathway impact score represented on the x-axis and −log_10_(p value) represented on the y-axis. Metabolomic analyses were performed in triplicate on samples.

To complement these transcriptomic insights, we performed metabolomic profiling using gas chromatography-mass spectrometry (GC-MS). Correlation analysis demonstrated good separation based on sample or metabolites (Figs. 4B and S1). We then proceeded to identify differentially regulated metabolites (Fig. 4C) and conducted pathway enrichment analysis (Fig. 4D-E). Of note, metabolites involved in Ala, Asp, and Glu metabolism were significantly decreased, which, combined with the METAFlux data, suggests that these metabolites have greater consumption following NUPR1 loss. Furthermore, glycolysis/gluconeogenesis metabolites were up-regulated while TCA cycle metabolites were down-regulated. These results are consistent with our findings of upregulated glycolytic gene expression as well as the multiple up- and down-regulated genes involved in the TCA cycle, suggesting these pathways are mis-regulated upon loss of NUPR1 in lung epithelial cells.

### NUPR1 is essential for mitochondria, regulation of oxidative stress, and tissue architecture in the mammalian lung

To determine the consequences of NUPR1 deletion *in vivo*, we evaluated *Nupr1*^-/-^ mice, which were grossly normal as previously reported (31). TEM of *Nupr1*^-/-^ lungs revealed a significant increase in both mitochondrial fission/fusion events (p=0.0485) and overall mitochondrial number (p=0.0060) compared with *Nupr1*^+/+^ lungs, consistent with our observation of increased mitochondrial content in epithelial cell lines (Fig. 5A-C). Furthermore, immunofluorescence for superoxide dismutase 2 (SOD2) and 8-OHdG demonstrated increased oxidative stress in *Nupr1*^-/-^ AT2 cells (Fig. 5D-E). Finally, H&E staining revealed airspace enlargement after exposure to CSE as quantified by cord length (Fig. 5F-G, p=0.0040). In summary, these results underscore the essential role of NUPR1 in regulating mitochondrial dynamics, modulating the oxidative stress response, and preserving tissue architecture in the mammalian lung.

**Figure 5.**
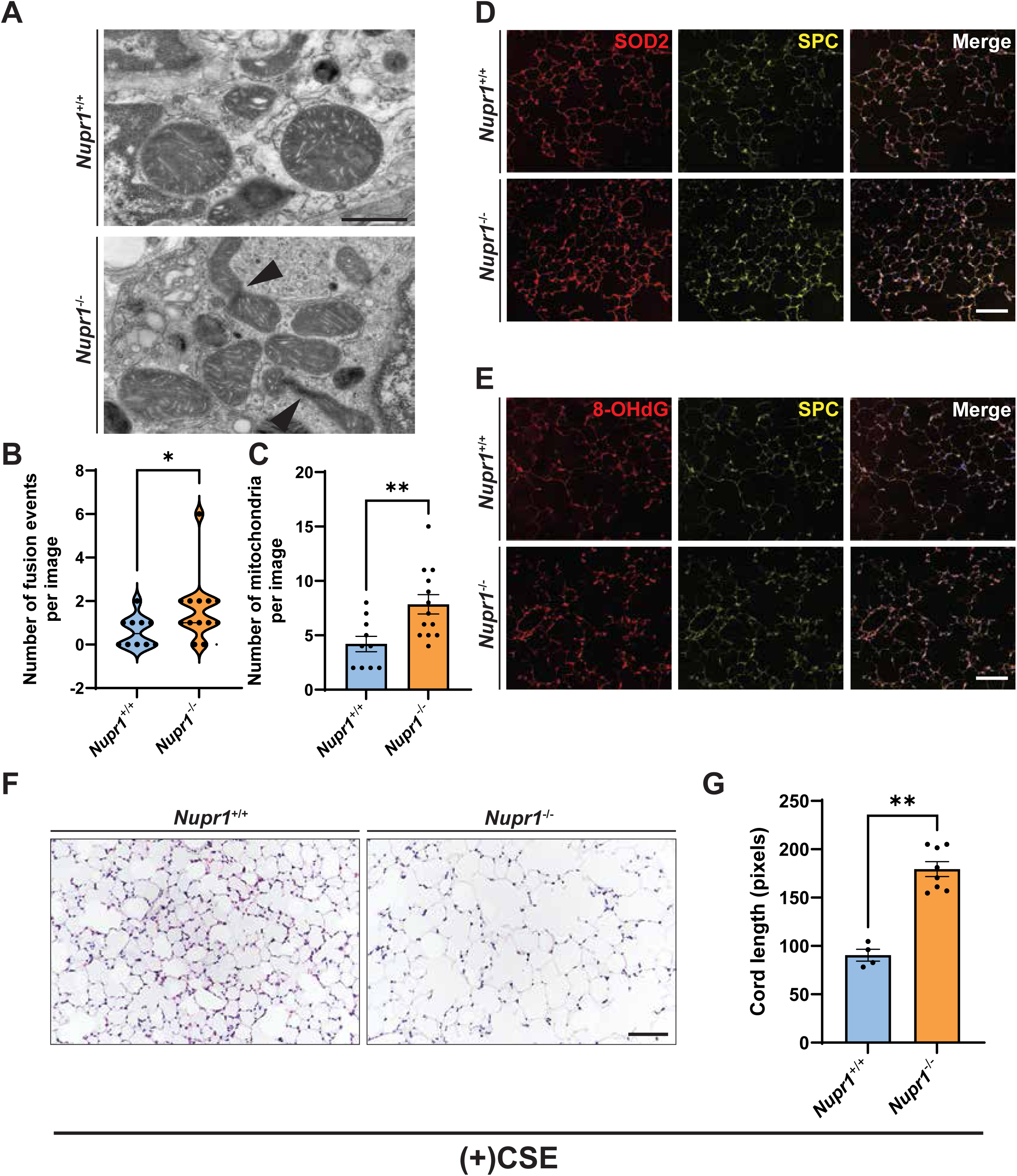
Loss of NUPR1 in mice causes defects in mitochondrial structure, cellular architecture, and pulmonary function in the lung. (A) TEM images of *Nupr1*^+/+^ and *Nupr1*^-/-^ lungs demonstrate increased mitochondrial fission/fusion events and increased numbers of mitochondria as quantified in (B) and (C), respectively. Arrowheads indicate fused mitochondria. Scale bar, 1 μm. Two-tailed T-tests were utilized to determine significance of changes in fission/fusion events and mitochondrial counted. Points indicate count obtained from an individual image. *, p<0.05; **, p<0.01. Confocal images reveal increased SOD2 (D) and 8-OHdG (E) staining (red) in AT2 cells (SPC, yellow) from *Nupr1*^-/-^ mouse lungs compared to *Nupr1*^+/+^ mouse lungs. DAPI is shown in blue in merged images. Scale bars, 100 μm. (F) H&E staining reveals increased airspace opacities in *Nupr1*^-/-^ mouse lungs compared to *Nupr1*^+/+^ mouse lungs exposed to CSE. Scale bar, 100 μm. (G) Quantification by mean chord length is shown with statistical analysis performed by two-tailed T-test. Points indicate count obtained from an individual image. **, p<0.01.

## Discussion

By employing functional, transcriptomic, and metabolomic experiments, we extend prior work and demonstrate that NUPR1 is an important regulator of cellular metabolism and bioenergetics, mitochondrial function, oxidative stress, iron metabolism, lipid peroxidation, and ferroptosis in the lung. Additionally, we translate these findings *in vivo* and show that lungs lacking NUPR1 demonstrate abnormal mitochondria, increased oxidative stress, and airspace enlargement after CSE exposure, suggestive of alveolar destruction. Given these findings and our previous work showing reduced NUPR1 expression in AT2 cells from patients with emphysema (28), our findings suggest dysregulated NUPR1 likely contributes to loss of alveolar homeostasis and disease progression in COPD.

NUPR1 is a stress response protein, acting as a transcriptional regulator transmitting multiple internal and external stress signals (22, 23). Recent studies identified NUPR1 as a ferroptosis inhibitor that regulates iron metabolism through an LCN2-dependent mechanism (25). In accordance with these observations, our data underscore the critical role of NUPR1 in ferroptosis regulation by uncovering its impact on iron metabolism, lipid metabolism, and the cellular redox state, all of which shape ferroptosis susceptibility. Notably, our transcriptomic analysis reveals increased *LCN2* expression following NUPR1 loss—a finding that contrasts with earlier reports of reduced LCN2 in pancreatic cancer cells upon NUPR1 depletion (25). This discrepancy suggests that the various functions of NUPR1, including the regulation of ferroptosis, may be highly context-dependent or cell-type specific.

NUPR1 loss has been associated with altered mitochondrial function (24, 27). Here, we extend these findings to the lung, demonstrating that NUPR1 depletion increases oxidative stress, reduces mitochondrial membrane potential, and impacts mitochondrial fission and fusion dynamics. Transcriptomic and metabolomic data indicate that loss of NUPR1 results in upregulation of glycolysis and simultaneously oxidative phosphorylation. However, an associated increase in adenosine triphosphate (ATP) was not observed in our metabolomic data, suggesting a decoupling of oxidative phosphorylation from ATP generation. Such a reprogramming likely redirects metabolites into other pathways (e.g., amino acid or fatty acid metabolism) rather than supporting efficient ATP generation via oxidative phosphorylation (39–41). Consistent with these observations, we found significant perturbations in the TCA cycle, glutamate/alanine/aspartate pathways, and the β-oxidation of lipids, all of which indicate a broader metabolic imbalance caused by NUPR1 deficiency.

Importantly, our as well as other recently published data raise the question to what extent does NUPR1 regulate alveolar epithelial plasticity in both healthy and diseased states (42, 43). In many developmental contexts, aerobic glycolysis favors a highly proliferative cell state associated with stemness whereas differentiation typically depends on oxidative phosphorylation (40, 44, 45). Consistent with this, previous work showed that deleting *Ndufs2*, a Complex I subunit, impairs the ability of AT2 cells to differentiate into alveolar type 1 (AT1) cells (46). These observations suggest that NUPR1 regulates the metabolic state of AT2 cells to influence their proliferative capacity and their ability to differentiate into AT1 cells. Additionally, as a cell stress response protein, NUPR1 is an important regulator of cellular adaptation whose expression can change depending on physiological and pathological conditions (21–23, 42, 43). In COPD, NUPR1 levels are reduced whereas, with age, NUPR1 expression tends to rise (unpublished data) (43). As a ferroptosis inhibitor (25), increasing NUPR1 should prevent ferroptosis-mediated cell death and promote cellular resilience. Furthermore, fluctuating NUPR1 levels influence stemness in and the ability of AT2 cells to transdifferentiate (42). Thus, by modulating NUPR1 levels, cells balance survival with the capacity for stem-like behavior and lineage shifts, underscoring its central role in regulating cellular fate.

Numerous limitations of this study and remaining critical questions deserve mention. Although we identified clear associations between NUPR1 deficiency and altered cellular metabolism and bioenergetics, further work is needed to determine how NUPR1 affects these processes on a mechanistic level. For example, the increased fission/fusion-like events we observe could reflect NUPR1-dependent modulation of core regulatory factors (e.g., DRP1, MFN1, MFN2, or OPA1) (47, 48). Alternatively, these changes may stem from a broader dysregulation of metabolic or stress-related pathways in the absence of NUPR1. Determining whether NUPR1 acts by directly controlling these key fission/fusion effectors or whether mitochondrial abnormalities emerge as a downstream consequence of global metabolic reprogramming will be essential for fully elucidating its role in maintaining mitochondrial homeostasis. Additionally, while we did not incorporate lipidomic analysis in this study, it remains an essential avenue for future exploration, given the reliance of surfactant production on lipid metabolism and the tight links between lipid metabolism and mitochondrial function (49–51). Beyond its downstream functions, an important question is how and why NUPR1 expression is specifically regulated in lung pathology, such as COPD. While epigenetic regulation has been proposed as one possible explanation (52, 53), more studies are needed to determine the exact mechanisms behind the cell-type and disease-specific expression patterns of NUPR1. Understanding these factors that control NUPR1 expression promises to further clarify its multifaceted role in lung disease and may ultimately lead to more targeted strategies for restoring alveolar homeostasis in COPD and related conditions.

## Data Availability

All datasets generated during this study will be made public and will be available upon request to corresponding author.

## Author Contributions

J.E.M. and M.S. conceived the study, T.Y., J.N., S-H.K., J.E.M., and M.S. designed experiments, T.Y., J.N., and S-H.K. performed the experiments, S.S.S., T.Y., J.N., S-J.K., and M.S. analyzed the data, and S.S.S. and M.S. wrote the manuscript. P. S-C. and J.I. assisted with experiments and provided critical reagents.

## Acknowledgements

The authors would like to thank past and present members of the Sauler lab for critical reading and input to this manuscript. Additionally, we thank the Metabolomics Core, Yale Center for Genome Analysis, and the Microscopy Core for assistance with metabolomics, transcriptomics, and TEM analysis, respectively.

## Supplementary Information

**Supplementary Figure S1.** Correlation analysis shows close correlation by metabolite for metabolomics data.

**Supplementary Table 1.** Table demonstrates Hypercarb column gradients. Mobile phase A was 15 mM ammonium formate, 0.03% acetyl acetone, and 0.1% formic acid while mobile phase B was 60% ACN, 35% IPA, 15 mM ammonium formate, and 0.1% formic acid.

**Supplementary Table 2.** Table shows Kinetex F5 column gradients. Mobile phase A was 95% water, 5% acetonitrile, and 0.1% formic acid while mobile phase B was 95% acetonitrile, 5% water, and 0.1% formic acid.

