## Supplementary figures and images for "The ferroptosis inhibitor NUPR1 coordinates the mitochondrial response to oxidative stress and cell metabolism during COPD pathogenesis in the lung"

### Supplementary Figure 1

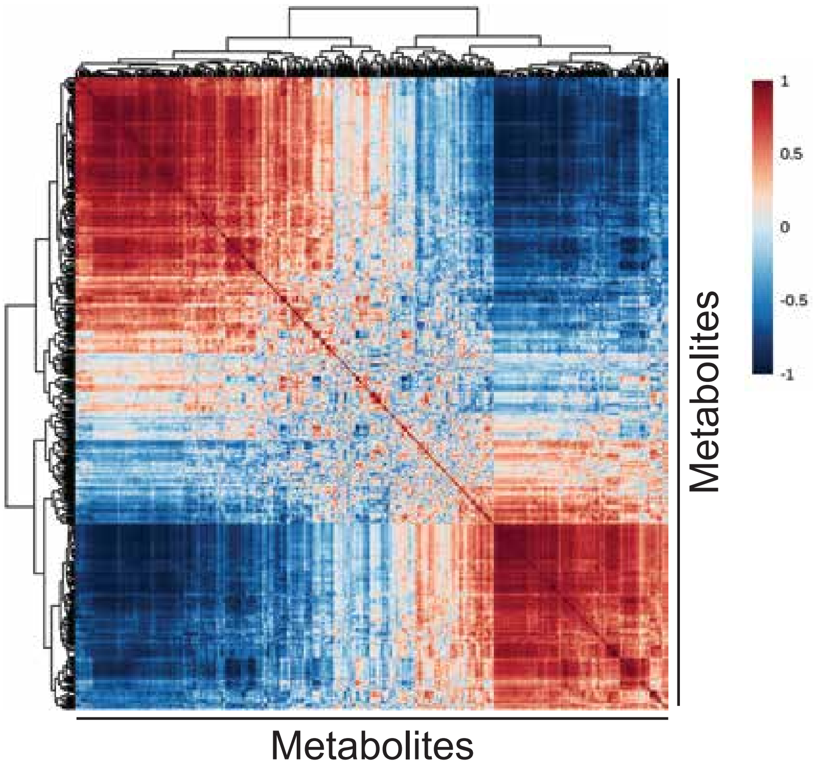


**Supplementary Figure 1**
