## Supplementary Table 1 for "The ferroptosis inhibitor NUPR1 coordinates the mitochondrial response to oxidative stress and cell metabolism during COPD pathogenesis in the lung"

| Time | Percent (%) of Mobile Phase A | Percent (%) of Mobile Phase B |
|---|---|---|
| 0 | 100 | 0 |
| 1 | 100 | 0 |
| 12 | 20 | 80 |
| 14 | 20 | 80 |
| 14.1 | 100 | 0 |
| 18.1 | 100 | 0 |

### Supplementary Table 1
