## Supplementary Table 2 for "The ferroptosis inhibitor NUPR1 coordinates the mitochondrial response to oxidative stress and cell metabolism during COPD pathogenesis in the lung"

| Time | Percent (%) of Mobile Phase A | Percent (%) of Mobile Phase B |
|---|---|---|
| 0 | 100 | 0 |
| 0.5 | 100 | 0 |
| 15 | 0 | 100 |
| 16 | 0 | 100 |
| 16.1 | 100 | 0 |
| 20 | 100 | 0 |

### Supplementary Table 2
